# Illuminating duckweed: hyperspectral reflectance highlights a predictive ceiling for accurate determination of biomass

**DOI:** 10.64898/2026.11.06.764931

**Authors:** Kellie E. Smith, Carlos A. Robles-Zazueta, Laura Cowan, Muhamad Hazwan Yahya, Lorna McAusland, Erik H. Murchie

## Abstract

Duckweed is a fast-growing aquatic plant with strong potential as a sustainable, high protein crop that can be produced without expanding agricultural land. However, substantial trait variation necessitates rapid selection proxies to identify genotypes with high growth, biomass and pigment content while simultaneously suited to low input, controlled environments. Although hyperspectral reflectance phenotyping is widely applied in conventional crops, its utility for duckweed remains unexplored. We assessed 133 UK accessions from six species grown under ambient light, quantifying growth, pigment composition, photosynthetic performance and spectral reflectance. A second panel with contrasting light tolerance strategies, including growth and photosynthetic acclimation, was used to test whether hyperspectral reflectance could detect photoprotective mechanisms associated with high light resilience. Duckweed accessions displayed extensive variation across all traits. However, vegetation indices (VIs) commonly used in arable crops demonstrated only limited predictive accuracy for total chlorophyll (R^2^ =0.34, p<0.005) and maximum photosystem II (PSII) quantum yield (*F*v/*F*m) (R^2^ =0.25, p<0.0001), with poor performance for other traits. Under high light, VIs moderately differentiated tolerant accessions, including those maintaining higher *F*v/*F*m, *Φ*PSII operating efficiency and carotenoid:chlorophyll ratios. Partial least squares regression (PLSR) modelling improved predictive performance, providing reliable estimates of chlorophyll content (R^2^ =0.5, RMSE =2.05 mg/g freeze dry weight) maximum non-photochemical quenching (NPQ) (R^2^ =0.5, RMSE =0.57) and NPQ efficiency of PSII (R^2^ =0.53, RMSE =0.04). Although predictions of biomass and growth rate remained weak, green and red edge spectral regions were strongly associated with pigment contents and photosynthesis and indirectly linked to biomass. This study provides the first comprehensive evaluation of spectral phenotyping for duckweed crop development. Our findings indicate that broad spectrum modelling approaches, integrated with other high throughput phenotyping tools, may support selection of high vigour varieties and monitoring of commercial duckweed production systems.

## 1. Introduction

The human population is predicted to increase to between 8.9 and 12.4 billion by 2100 (United Nations; 2022). However, current food systems are insufficient to support this population growth, mainly due to limited availability of land, soil depletion and water scarcity (FAO; 2021). Alternative sources of food which can be produced with lower environmental footprints could contribute to improved food security. To this end, duckweed are miniature freshwater plants consisting of more than thirty six species with worldwide distributions. Under the name of ‘water lentils’ they have been earmarked as a new sustainable crop due to fast asexual reproductive rates (Sree *et al*.; 2015b, Michael *et al*.; 2021), high protein compositions (Leng *et al*.; 1995, Ziec *et al*.; 2025), high macronutrients and carotenoids content (Appenroth *et al*.; 2017, 2018, Xu *et al*.; 2023) and ability to grow in controlled hydroponic systems, including for space agriculture (Coughlan *et al*.; 2022, Petersen *et al*.; 2022b, Polutchko *et al*.; 2022).

Duckweed species and varieties thereof have displayed different growth rates in these systems (Ziegler *et al*.; 2015) with contrasting adaptation strategies to high light environments (Smith *et al*.; 2024a, Xuan *et al*.; 2025), and unsurprisingly different nutrient, protein, amino acids, carotenoids and secondary metabolite profiles between species and controlled environments (Landolt, 1986; McCann, 2016, Stewart *et al*.; 2020, 2021, Smith *et al*.; 2024a, 2024b, 2024c, 2025). Growing duckweed for maximal production of fresh biomass and freeze-dried biomass in low input systems are optimal target traits for food production. For example, production of fresh biomass could be used as a vegetable source and freeze-dried biomass could be used to produce plant-based protein supplement powders. Concordantly, duckweeds also produce numerous protection compounds known to be beneficial when included in human diets. For instance, while zeaxanthin is a predominant plant carotenoid formed as a protective mechanism from de-epoxidation in high light, its consumption supports eye health, the immune system and it has shown to enhance anti-tumour immune responses (Zhang *et al*.; 2025). Consumption of plant chlorophylls and other secondary metabolites can provide high sources of amino acids, macro- and micronutrients. Therefore, maximization of duckweed growth to produce high biomass containing these key food quality traits using respective light and nutrient environments are of interest to growers (Petersen *et al*.; 2022a). To meet their economic potential as a protein source or vegetable, strategies to select optimal duckweed candidates offering these traits should be developed by phenotyping in suitable growth environments.

To develop duckweed with perspectives towards alternative food sources, it is vital to select for performance in response to environment and differentiate the varieties based on their vigour in low input systems. Pigment production directly relates to light capture and influences growth and stress responses in plants. . Estimates of ‘greenness’ or leaf chlorophyll content from spectroradiometers are commonly used as health indicators and nitrogen (N) status of staple field crops, e.g. maize (Shivashankar *et al*.; 2025), wheat (Islam *et al*.; 2014) and rice (Kim *et al*.; 2024). This trait is especially important for the selection of leafy edible crops like spinach, lettuce or indeed duckweed (Gray *et al*.; 2026). Unfortunately, it is challenging to use leaf clamp-on methods in duckweed due to small sizes of fronds (instead of leaves) together with their aquatic growth habits. Other existing quantitative measurement of duckweed health or ‘greenness’ require laboratory-based analysis using time- consuming destructive methods including chlorophyll extraction (Calzadilla *et al*.; 2022). Due to their small sizes, total chlorophyll content is more easily achieved in duckweeds as a per unit biomass rather than per area basis as applied in broader leaved crop plants. Chlorophyll content in duckweed depends on the different species adaptation to their light environment and growth strategies (Strzalek & Kufel.; 2021). While chlorophyll content has been reliable to study acclimation in small number of accessions, it remains a slow approach for phenotyping large populations (Smith *et al*.; 2024a). Therefore remote approaches must be sought for phenotyping of green pigments to select for high vigour varieties and to monitor health of duckweed populations.

Plant vigour is closely linked to photosynthetic capacity and light use efficiency, making these traits particularly important in duckweed, the world’s fastest-growing flowering plant. Understanding the physiological basis of variation in light capture and growth is essential for optimizing biomass accumulation, productivity and micronutrient production e.g. carotenoids for food production. Gas analysers and chlorophyll fluorescence have the utility to screen plants for enhanced photosynthetic performance and can predict yield in challenging environments (Pavlovic *et al*.; 2014, Carmo-Silva *et al*.; 2017). However, these techniques require expensive specialized equipment and have typically lagged behind research in traditional arable crops. Chlorophyll fluorescence provides a rapid, non- destructive measure of photosynthetic performance and has proven useful for assessing plant health in duckweed (Stewart *et al*.; 2020, 2021). Key traits include maximum quantum yield (*F*v/*F*m), which reflects the maximum efficiency of photosystem II (PSII); PSII operating efficiency (ΦPSII) and electron transport rate (ETR), which measure the efficiency of photochemistry and electron flow under a given light environment; and photoprotective processes including non-photochemical quenching (NPQ) and regulated thermal dissipation (Y(NPQ)), which dissipate excess absorbed light energy as heat, largely through the xanthophyll cycle. Together, these traits provide insight into the balance between photosynthesis and photoprotection that underpins growth and adaptation to contrasting light environments. Complementary measurements of carbon assimilation have also recently become possible in duckweed through adaptations of closed-system gas exchange analysers (Rintoul *et al*.; 2026). Although still in its infancy, the adaptation of high throughput techniques such as hyperspectral reflectance offers a promising non-invasive phenotyping approach for large duckweed germplasm

collections. The application to duckweed could enable the identification of trait proxies associated with photosynthetic capacity, pigment composition, photoprotection and growth-related traits or ‘vigour’ in the context of controlled high and low growth environments designed for food production (Sanaeifar *et al*.; 2022).

Hyperspectral reflectance signatures, which can be obtained across simultaneous wavelengths in a single scan or image, has proved useful in optimizing crop nutrition (Pandey *et al*.; 2017, Mahajan *et al*.; 2017), predicting photosynthetic capacity (Robles-Zazueta *et al*.; 2021; 2022, Zhi *et al*.; 2022), disease detection (Shi *et al*.; 2018), for cultivar classification (Lyu *et al*.; 2024) and maximizing crop yield (Montesinos-López *et al*.; 2017). These approaches have contributed to the improvement of conventional crops using breeding panels. However, these approaches have not been explored in the novel underutilised crop duckweed to select natural varieties which already feature enhanced nutrition, photosynthesis or growth performance in high or low controlled environments. Whilst, breeding is challenging to achieve in broadly asexually reproducing duckweed, selecting varieties by phenotyping proxy traits could aid their selection for upscaled production. Furthermore, hyperspectral sensing could become a tool for monitoring health and productivity of scaled-up operations in duckweed systems or bioreactors.

Although hyperspectral phenotyping is extensively used in plant breeding, ecophysiology and taxonomy studies, its use in duckweed remains currently limited. For example, leaf spectra from the duckweed species *Lemna gibba* was used to detect stress responses to a chemical contaminant through changes in pigment absorption features (Dong *et al*.; 2002). More recently, multispectral imaging was combined with chlorophyll fluorescence to assess duckweed tolerance to high elemental loads through changes in *F*v/*F*m, supporting the selection of varieties for water remediation (Audenaert *et al*.; 2026). In a food security context, higher duckweed tissue glucose and amino acid contents was linked with higher green reflectance vegetation indices derived from hyperspectral reflectance of growing duckweed populations (Smith *et al*.; 2024c). However, there currently remains unexplored opportunities to use hyperspectral information for selection of food production traits such as biomass production, nuanced light harvesting components of photosynthesis and protective pigment production in the prospective vegetable crop duckweed.

Despite the development of methods for interpretation of hyperspectral data, limited prediction testing has been used for diverse traits in duckweed. Vegetation indices (VIs) are computationally efficient and readily interpretable mathematical operations developed for arable crops. Vegetation indicies target key spectral signatures associated with specific traits but potentially overlook subtle genotypic biochemical and biophysical variation captured elsewhere in the spectrum. By contrast, analysis of contiguous narrowband hyperspectral spectra preserves detailed spectral information, enabling the detection of absorption features linked to pigment composition, nutrient and water status, and stress-induced biochemical, biophysical and physiological responses (Gamon *et al*.; 2019). Careful consideration of analytical approaches may be particularly important for aquatic plants, such as duckweed, which exhibit rapid metabolic and morphological adaptations to their environment. In addition, the use of hyperspectral reflectance data to predict biomass accumulation, light harvesting traits and growth in aquatic macrophytes has already been shown to improve trait accuracy using whole spectra compared to VIs (Villa *et al*.; 2017, Stratoulias *et al*.; 2015). Therefore, proxies for accurate trait prediction could be employed as part of variety selection and monitoring performance in controlled environments used for duckweed food production.

Hyperspectral linkage between biomass productivity and light resilience traits within and between duckweed species has not been assessed. The objective of this study was to investigate variation in vigour of a cohort of 133 UK duckweed varieties from six major species in common controlled environments. We evaluated the predictive ability of vegetation indices (VIs) and hyperspectral signatures spanning the visible and near-infrared (NIR) spectrum (400-1100 nm) using multivariate modelling approaches. To achieve this, different proxies of plant productivity were measured in the laboratory including chlorophyll fluorescence dynamics, pigment composition and plant growth rates primarily in ambient low growth light (GL), with a smaller cohort in a high growth light (GL) environment. Our hypothesis is that reflectance wavelengths in the green and red regions of the electromagnetic spectrum should positively associate with chlorophyll content, while NIR regions will be linked with photosynthetic performance and higher growth rates.

## 2. Materials and methods

### 2.1. Duckweed collection

One-hundred and thirty-three accessions of six species described in (Smith *et al*.; 2025) were used as a duckweed study panel to test the utility of hyperspectral reflectance to predict growth rate, photosynthetic traits and pigment content as relevant traits for selection of novel food crops. The selected species include: *Spirodela polyrhiza*, *Lemna minor*, *Lemna japonica*, *Lemna gibba*, *Lemna minuta* and putative hybrid species *Lmu/Lmo* (*L. minor* x *L. minuta*). The UK duckweed cohort was chosen for their species-level classifications, associated characterisation of their historical native pond light environments and recent domestication to controlled environments (summarized in Table S1). The data collection pipeline to assess relevant food crop traits in this study is presented in Fig. 1.

**Figure 1.**
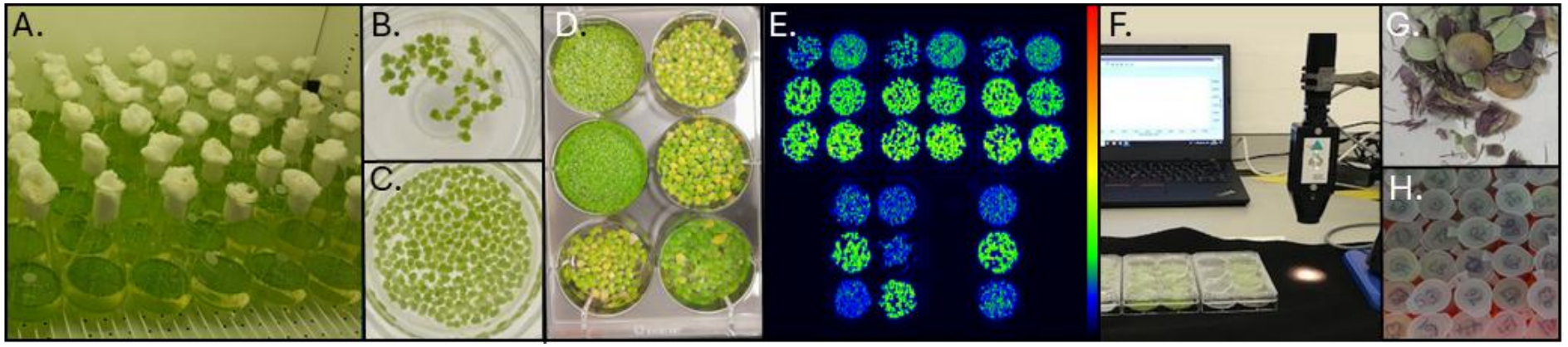
Workflow for characterisation of duckweed accessions towards novel food development. **A.** Duckweed accessions were grown in individual flasks in controlled environments supplying low growth light (GL) conditions (50 µmol m^-2^ s^-1^). **B-C**. Growth rate of biomass in flasks was assessed weekly for four weeks. **D.** Duckweed biomass was split into three replicates in well plates after each growth experiment. **E.** Duckweed biomass was imaged for chlorophyll fluorescence after dark adaptation to produce false color image. Measurements were conducted for each replicate population using an imaging system (800C Fluorcam, Photon System Instruments, Brno, Czech Republic). **F.** Hyperspectral reflectance was measured from duckweed biomass with a spectroradiometer (ASD FieldSpec 4, Malvern Panalytical, UK). **G.** Total fresh biomass for each accession was weighed and freeze-dried. **H.** Freeze-dried biomass was aliquoted into known mass for destructive pigment extractions using acetone to obtain total chlorophyll and carotenoid contents per mass. Each accession from low GL experiments (*n* = 133) was evaluated across three independent growth rate experiments. Per accession, this resulted in nine chlorophyll fluorescence and nine hyperspectral datasets (three per growth rate experiment each) and twelve chlorophyll extraction measurements (four per growth rate experiment). For the high GL experiment, 24 accessions were each subjected to five growth rate experiments, resulting in fifteen chlorophyll fluorescence datasets and fifteen hyperspectral datasets (three per experiment), and twenty chlorophyll extraction measurements (four per experiment) per accession.

### 2.2. Duckweed growth experiments

Duckweed accessions were grown in low (50 µmol m^-2^ s^-1^) GL experimental conditions for four to six weeks (Fig. 1A-C). GEN-1000 controlled environment cabinets (Conviron, Winnipeg, Canada) provided broad-spectrum LED lights giving a light photoperiod of 16:8 hr days and temperature cycle of 22 °C in the day and 18 °C during nights. Duckweed varieties were grown separately in Erlenmeyer flasks for each growth rate experiment as batches according to the UK regions they were collected. At the start of each experiment, a single colony was used to initiate the population and growth was then measured weekly for each accession using (i). changes in green area normalized to initial frond area (T0) using RGB from photos and (ii). colony counts. N-medium was made (as described in Appenroth *et al*.; 1996) and refreshed weekly to provide plentiful nutrient conditions for growth. Relative growth rate (RGR) was obtained from total green area [Eq1] and from colony number [Eq2] during the log phase of growth. After four weeks, the fastest growing accessions were harvested as they had exhausted the surface growing space. Slower growing accessions were grown for a further two weeks with continued N-medium replenishment to expand biomass yield to ∼95% surface coverage. Each duckweed growth experiment was repeated as batches for the 133 accessions three times over a three- year period to obtain three biological replicates per accession.

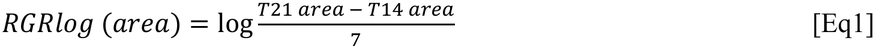

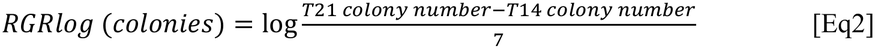

Adaptation to controlled high GL was measured in a small subset of 24 accessions described in (Smith *et al*.; 2024a, highlighted in Table S1). *Lemna gibba* and the putative hybrid species *Lmu/Lmo* was not included in this regional cohort. Instead, the parent species *Lemna turionifera* which forms the hybrid *L. japonica* (*L. minor* x *L. turionifera*) featured exclusively in this specific region (Smith *et al*.; 2025) These duckweed accessions were selected for comprising species diversity and representing a local cohort with spatial and seasonal light variability in their water habitats. In addition to their growth in low GL environments, these accessions were grown concurrently under high GL conditions (350 µmol m^–2^ s^–1^) with all other environmental parameters kept consistent and growth rates measured as per the low GL experiments. High GL experiments on this subset were repeated five times over the three-year period to give *n*=5 replicates. During each GL experiment, the light environment (photosynthetically active radiation; PAR, light intensity and quality) for each duckweed population was measured in the controlled environments using a light meter (Li-250A, Licor Biosciences, NE, USA) and handheld spectrometer (LI-180, Licor Biosciences, NE, USA) in each growing position to account for spatial variation.

### 2.3. Chlorophyll fluorescence as an indicator of duckweed photosynthetic health

At the end of growth experiments, each accession represented by a duckweed population in each flask was subset three times into separate six well plates to obtain a total of nine and fifteen biological replicates for the low and high GL conditions respectively for chlorophyll fluorescence measurements. Each well contained 3 ml N-medium and the surface of each well was filled (where possible) with dense populations of each accession to maximize biomass signal and minimize duckweed colony repellence and water visibility in each well (Fig. 1D-E). A closed chlorophyll fluorescence imager (800C Fluorcam, Photon System Instruments, Brno, Czech Republic) was used to measure photosystem II (PSII) chlorophyll fluorescence light reactions for each accession after 1 hr dark adaptation. A light response curve protocol featuring eleven incremental light levels was setup as described in (Smith *et al*.; 2024a). Stepwise light levels were calculated to corresponding photosynthetic photon flux density (PPFD) levels to select three contrasting light levels to capture specific light adaptive responses after each GL condition. This included the light level corresponding closest to the low GL environment (LL) (130 µmol m^-2^ s^-1^), high GL environment (HL) (365 µmol m^-2^ s^-1^) and very high light (VHL) (>1000 µmol m^-2^ s^-1^), which was the maximum light level experienced during the light curve but not experienced in either GL environment. Photosynthetic traits from chlorophyll fluorescence were calculated as outlined in (Murchie & Lawson, 2013). Maximum quantum yield of PSII (*F_v_*/*F_m_*) in the dark was calculated using [Eq3], operating PSII operating efficiency (ΦPSII) was calculated as in [Eq4] and non-photochemical quenching (NPQ) as in [Eq5] for each light level.

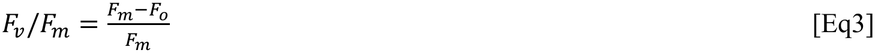

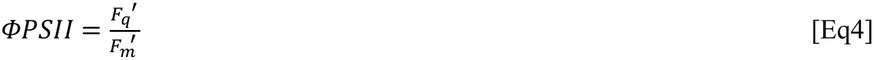

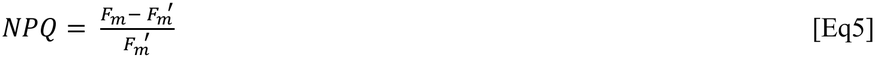

In addition to these, the following traits were derived from (Maxwell & Johnson; 2000, Kramer *et al*.; 2024): Electron transport rate (ETR) at LL (130 µmol m^-2^ s^-1^) was calculated using incident PAR and ΦPSII [Eq4] as in [Eq6]. Constants for this equation included 0.84 as the default fraction of incident light absorbed by a leaf and 0.5 corresponding to average ratio of PSII:PSI reaction centres. Then maximum NPQ at VHL (1050 µmol m^-2^ s^-1^ VHL) was used to derive the yield of PSII regulated non- photochemical energy dissipation as heat (Y(NPQ)) using [Eq7].

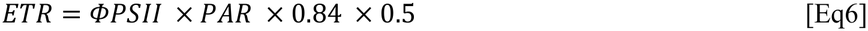

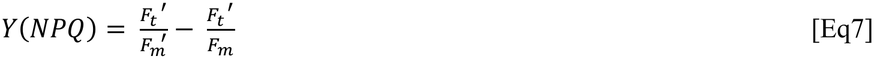

### 2.4. Hyperspectral signatures from duckweed biomass

Immediately after growth in low or high GL, followed by chlorophyll fluorescence experiments, a spectroradiometer (ASD FieldSpec 4, Malvern Panalytical, UK) was used to measure the reflectance signatures of duckweed biomass setup in six well plates (Fig. 1F). Reflectance was measured from adaxial frond surfaces at 1 nm increments between 350 – 2500 nm in a laboratory providing consistent low light at 5 µmol m^-2^ s^-1^. A contact probe attachment with a halogen bulb light was suspended in a tripod placed at a constant distance of 20 cm directly above each duckweed accession population. A black background was placed behind well plates to reduce stray reflectance e.g. from N-medium and metallic lab bench. A white reference panel was scanned every five minutes to re-calibrate the instrument between measurements. Three pseudo-replicate measurements were taken per well to capture hyperspectral variation within well populations, and biomass was placed in three individual plates, giving a total of nine biological replicates for each accession (*n* = 9) after each growth rate experiment.

### 2.5. Harvesting duckweed biomass and destructive chlorophyll extraction

After hyperspectral measurements, the three replicates per accession were removed from the well plates, air dried for 20 mins and weighed to obtain the total fresh biomass weight from each growth rate experiment. These replicate populations were then frozen in liquid nitrogen, freeze-dried for 2 days and then re-weighed to obtain freeze-dried biomass for each accession replicate (Fig. 1G). Each of these replicates were portioned into 5 mg whole tissue samples for pigment extraction (Fig. 1H). Chlorophyll and carotenoids were extracted by acetone to quantify pigments on a per mass basis and detected by absorption using spectrophotometry as described in (Smith *et al*.; 2024a).

### 2.6. Spectral data analysis

Raw reflectance signatures from lab-grown duckweed were pre-processed using the signal package in R (Ligges *et al*; 2015) with a Savitzky–Golay filter to smooth and reduce noise. Two analytical approaches were then applied. First, reflectance measured at specific regions of the electromagnetic spectrum was used to calculate vegetation indices (VI), which are derived from ratios and differences of peaks in the visible, near-infrared (NIR) and shortwave infrared regions (Xue & Su.; 2017). Commonly used VIs were derived from blue, green, red, red edge and NIR spectral regions. Five VIs (NDVI, GM, PRI, OVI, EVI) commonly used for prediction of vigour traits in plants and crops were selected from the literature (Table 1) and calculated for each accession replicate under low GL conditions and with the subset panel in high GL conditions. Relationships between VIs and physiological traits were assessed using Pearson’s pairwise correlations using the *cor* function from the stats package and visualized using a matrix from the corr package (Wei & Simko.; 2024). Then key associations were modelled for linear mixed-effects models fitted with *lme4* (Bates et al.; 2015). Prior to modelling, alternative functional forms (linear, logarithmic, square-root, polynomial, nonlinear) were evaluated using Akaike Information Criterion and residual diagnostics and traits exhibiting non-normality or heteroscedasticity were transformed. Models included light intensity PAR measured in each growth position as a fixed effect and accession as a random intercept, with species identity incorporated either as a random effect or as a fixed interaction. Model performance was quantified using marginal R² (fixed effects) and conditional R² (fixed + random effects).

**Table 1.** Common hyperspectral vegetation indices (VI) derived from (Index Database, 2026) and their main uses to predict plant traits including crop yield and chlorophyll content. *R* = Reflectance at wavelength.

| VI | Full name | Formulae | Estimates | Plants | References |
|---|---|---|---|---|---|
| NDVI | Normalized difference vegetation index | $\frac{R800-R680}{R800+R680}$ | Green biomass, leaf area index, chlorophyll content | 13 spp.<br><50 spp.<br>Rice<br>Sorghum | Gamon *et al*.; 1995<br>Sims & Gamon; 2002<br>Shi *et al*.; 2018<br>Zhang *et al*.; 2022 |
| EVI | Enhanced vegetation index | $2.5*\left(\frac{R800-R670}{R800+6*670-7.5*R470+1}\right)$ | Crop yield | Soybean<br>Rice | Shammi & Meng; 2021<br>Son *et al*.; 2014 |
| GM | Green model | $\frac{R750}{R550}-1$ | Chlorophyll content | 6 spp.<br>Citrus | Gitelson & Merzlyak; 1997<br>Xiao *et al*.; 2024 |
|  |  |  | Glucose, amino acid content | Duckweed (*S. polyrhiza, Lemna* spp) | Smith *et al*.; 2024c |
| PRI | Photochemical reflectance index | $\frac{R531-R570}{R531+R570}$ | Photosynthesis, carotenoids, xanthophyll cycle | 20 spp.<br>>100 spp. | Gamon *et al*.; 1997; Garbulksy *et al*.; 2011 |
| OVI | Optimal vegetation index | $R760-R730$ | Nitrogen | Wheat | Erdle *et al*.; 2011 |
|  |  |  | Fresh biomass | Duckweed (*S. polyrhiza, Lemna* spp.) | Smith *et al*.; 2024c |

For the second approach, partial least squares regression (PLSR) was used to model the association between pigments, photosynthetic and biomass traits with hyperspectral reflectance signatures. The spectral range used for PLSR was 400-1100 nm to reduce the water absorption bands in the shortwave infrared range (>1200 nm). PLSR is a supervised regression approach that maximizes the covariance between highly dimensional and collinear predictors (i.e. spectral reflectance at thousands of wavelengths) and the response variable. Trait predictions were done using the orthogonal scores PLS algorithm from the R *pls* package (Mevik & Wehrens.; 2007). The hyperspectral datasets were split in 80% for model training and 20% for model validation by species. The optimum number of components was determined through cross-validation at the training stage to select the model with lower root mean square error (RMSE). Finally, model performance metrics such as the determination coefficient (R^2^), RMSE and model bias were used to select the best model at the validation stage (Robles-Zazueta *et al*.; 2022).

## 3. Results

### 3.1. Varied performance of duckweed cohort for crop- associated traits

The duckweed cohort showed marked phenotypic variation in pigment contents, photosynthesis, photoprotection, growth and biomass trait groups in a common low GL controlled environment (Fig. 2). The clear phenotypic separation among accessions highlights their value as a diverse panel for evaluating hyperspectral signatures as a selection tool. Briefly, *Lemna japonica* accessions demonstrated consistently higher chlorophyll and carotenoid contents and higher fresh and freeze-dried biomass, linking pigment content to productivity (Fig. 2A, C). Northern *Lemna minor* accessions displayed high photoprotective capacity under ambient light, achieving the highest maximum NPQ and greatest regulatory dissipation as heat as Y(NPQ) (Fig. 2B). NPQ at very high light (VHL) significantly varied between species and geographic origins, with accessions from the highest latitudes showing the strongest NPQ responses (p <0.0001) but yielding less biomass. This significant variability in vigour and trade-offs due to geographic origin demonstrates that not all accessions have idealized crop traits in low input environments.

**Fig. 2.**
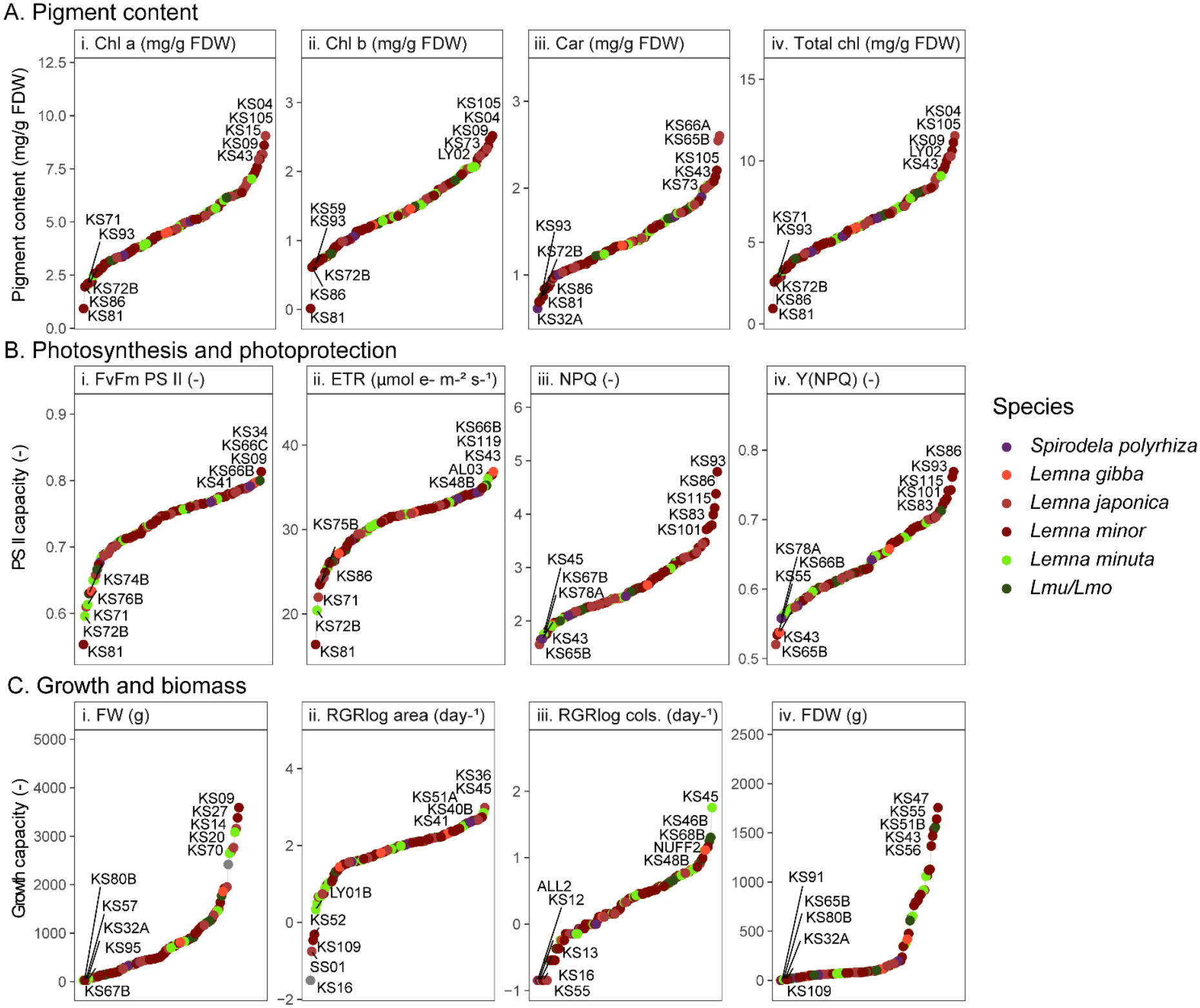
Varied performance of 133 duckweed accessions in low GL environments for crop-associated traits. **A.** Pigment contents measured by pigment extraction and spectrophotometry in mg g^-1^ freeze- dried weight: i. Chlorophyll a, ii. Chlorophyll b, iii. Carotenoids and iv. Total chlorophyll (a+b). **B.** Photosynthesis and photoprotection measured by chlorophyll fluorescence: i. Maximum quantum yield of PSII measured after dark adaption (*F*v/*F*m), ii. Electron transport rate (ETR) measured at LL (130 µmol m^-2^ s^-1^), iii. Non-photochemical quenching (NPQ) at VHL (1050 µmol m^–2^ s^–1^) and iv. Regulated thermal dissipation (Y(NPQ)) measured at VHL. **C.** Growth and biomass traits: i. Fresh weight (FW) biomass (g) at the end of growth period, ii. RGRlog area measured by green area increase during log phase, iii. RGRlog colonies based on colony count gain during log phase and iv. freeze-dried weight (FDW) at the end of the growth period. Unitless traits are indicated with (-).Trait averages were used from all replicates for each accession, the top and bottom five accessions (based on accession averages) are labelled and points for each accession are coloured by species.

### 3.2. Evaluating vegetation indices as predictors of pigment composition, photosynthetic performance and growth

Several positive correlations were observed between measured pigment contents (chlorophyll a, b and total chlorophyll), electron transport rate and VIs derived from the green and red spectral regions, particularly the GM index and OVI which incorporates the NIR region (Fig. 3A, 3B). Pigment concentrations, photosynthetic traits and VIs including GM, NDVI and OVI showed strong collinearity, indicating overlaps in trait and spectral regions attributed to vigour. Vegetation indices which produced the highest R^2^ for key selected traits are summarized in Fig. 3. The strongest relationship was between total chlorophyll and GM: accessions with higher chlorophyll content consistently showed higher GM values (Fig. 3A). While maximum quantum yield (*F*v/*F*m) was most strongly related to OVI this was a weaker correlation, with satisfactory separation of high and low performing accession clusters on the axes (Fig. 3B). Model fits improved substantially when accession-level effects were included (chlorophyll: R² = 0.337 to 0.501; *F*v/*F*m: R² = 0.250 to 0.432), indicating important accession and species-specific variation in these traits. Species-specific responses influenced GM predictions for total chlorophyll and OVI predictions for quantum yields, indicating species-specific differences in these traits under low GL conditions.

**Fig. 3.**
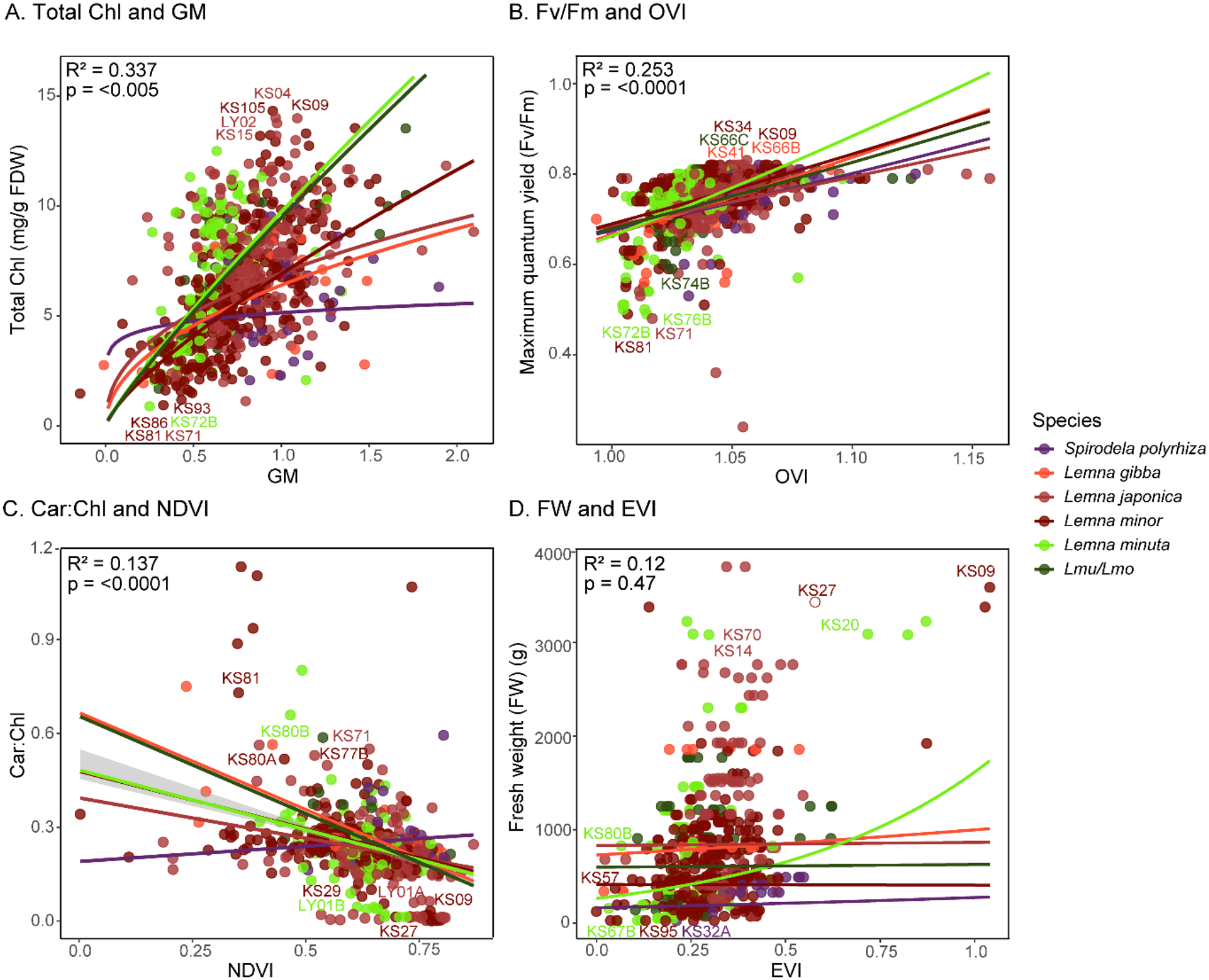
Associations between pigment concentrations, photosynthesis and fresh biomass with selected vegetation indices (VIs) using 133 duckweed accessions from a low GL environment. **A.** Total chlorophyll concentration vs GM. **B.** Maximum quantum yield (*F*v/*F*m) vs OVI. **C.** Carotenoid:chlorophyll ratio vs NDVI. **D.** Fresh weight (FW) biomass vs EVI. Mixed-effects regression models were fitted for each trait–VI relationship.Log transformation was used for pigment content and fresh weight; square-root transformation was used for *F*v/*F*m. Fitted slope lines show model predictions for each species. Marginal R² values and model p values are reported for each panel. Points represent averages for each accession from experimental replicates and are coloured by species type.. The top and bottom five accessions for each vigour trait are labelled.

Although several VIs showed statistically significant associations with physiological traits, marginal R² values < 0.5 indicated limited prediction power of VIs in low GL conditions.NDVI showed a weak, negative relationship with carotenoid:chlorophyll ratio, with substantial overlap among accessions preventing clear separation of high and low trait values (Fig. 3C). The model explained just 13.7% of trait variation, increasing when accession effects were included (conditional R² = 0.416), indicating substantial accession- and species-level variation not captured by NDVI.

Total fresh weight (FW) biomass was a key trait of interest, given the substantial variation found between accessions (Fig. 2Ci) and requirement for food production. NDVI showed little association with FW despite its ability to predict yield in arable crops, so EVI was evaluated as an alternative index to account for canopy density and reduce saturation in the red edge region between red and NIR. Although EVI showed a weak, positive association between with FW, high and low biomass-producing accessions overlapped extensively. The mixed effects model using EVI explained only 12% of variance and EVI was not a significant predictor overall (p = 0.47), indicating that VIs using NIR tested here have limited potential to predict biomass production in dense duckweed conditions (Fig. 3D).

Despite clear variation in photoprotective traits among accessions (Fig. 2B), including carotenoid content, xanthophyll cycle activity via NPQ and regulated heat dissipation as Y(NPQ), no VI showed a strong association with these traits under low GL conditions. This contrasts with the established use of PRI to predict photoprotective responses in arable crops. To test whether spectral sensitivity improves when photoprotection is more strongly induced, a subset of accessions with contrasting light adaptation strategies was exposed to high GL.

### 3.3. Association between VIs and high light stress responses

A subset of twenty-four accessions selected from (Smith *et al*.; 2024a, 2025) was grown under high GL conditions and VIs derived from hyperspectral reflectance were evaluated for their ability to predict light adaptive traits. As in the low GL experiment, several positive correlations emerged, most notably between maximum quantum yield *F*v/*F*m, operating efficiency (*Φ*PSII) measured at three key light levels (LL, HL, VHL) and predictor indices from the green, red and NIR spectral regions, particularly PRI and NDVI (Fig. 4A). However, even under high GL when photoprotection is induced, no VI showed a significant association with NPQ measured at any light level (LL, HL, VHL). This indicates that the VIs still lacked sensitivity to capture photoprotective responses in duckweed grown under high GL environments.

**Fig. 4.**
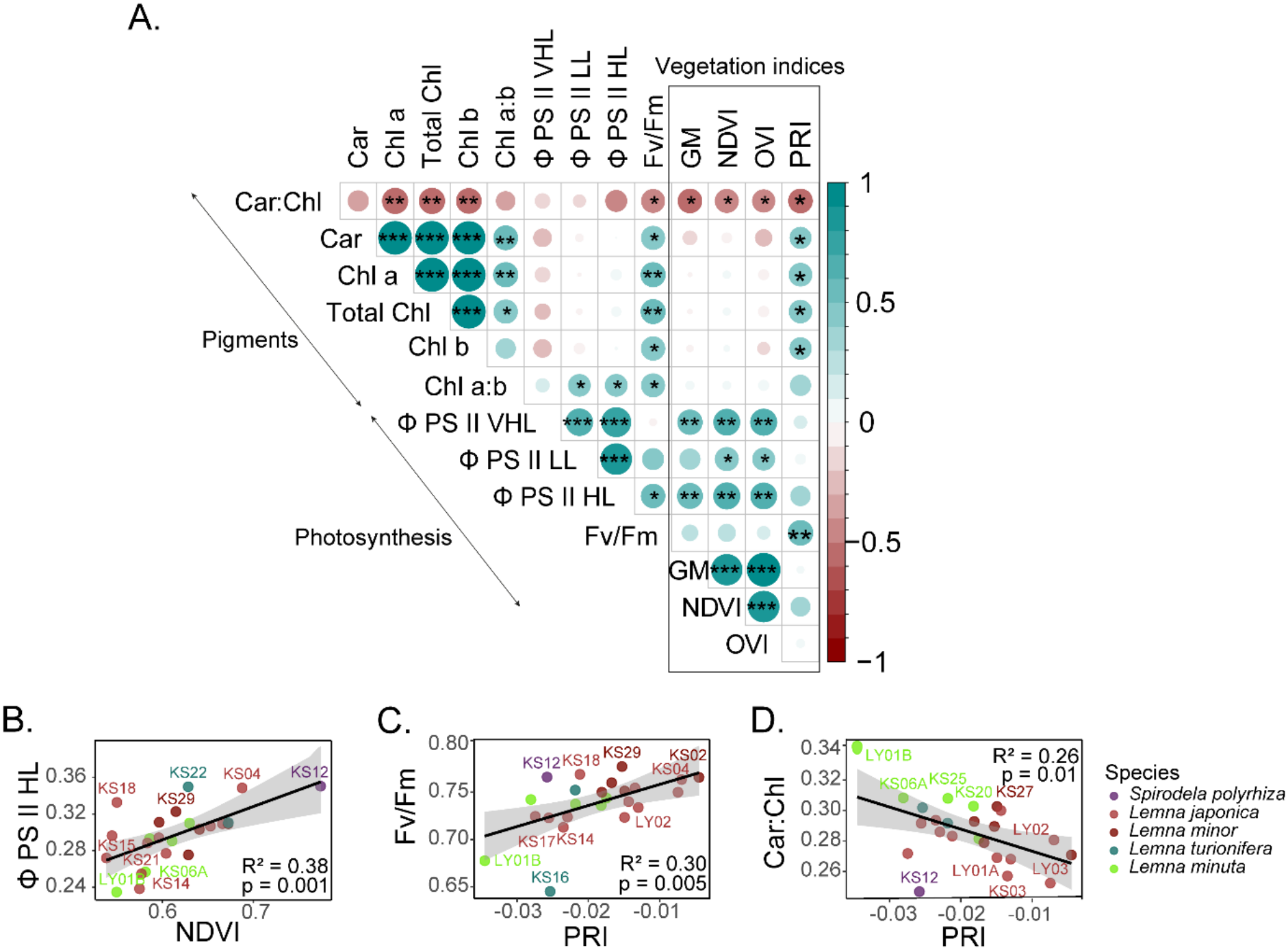
Significant associations between VIs with pigment and photosynthetic traits in a subset of 24 duckweed accessions from high GL experiment. **A.** Pearson correlation matrix showing relationships among pigment traits, photosynthetic traits and reflectance-derived VIs. Vegetation indices (GM, NDVI, PRI, OVI) are outlined by a box. Color and scale indicate direction of association, circle size indicates strength of association and asterisks denote statistical significance (*** p <0.001, ** p <0.01, * p <0.05). **B-D.** Pairwise relationships between measured traits and predictive VIs. **B.** *Φ*PSII operating efficiency at HL (365 µmol m^-2^ s^-1^ ) association vs NDVI, **C.** Maximum quantum yield (*F*v/*F*m) vs PRI, **D.** Carotenoid:chlorophyll ratio vs PRI. Points represent averages for each accession from replicates and are colored by species. The top and bottom five accessions for each measured trait are labelled.

Multiple VIs (GM, NDVI, OVI, PRI) showed negative associations with carotenoid:chlorophyll ratios (Fig. 4A, D), mimicking the trends found in the low GL experiment with the larger cohort (e.g. Fig. 3C). In the high GL condition, NDVI was positively associated with *Φ*PSII operating efficiency across all three key light levels (Fig. 4A), with the strongest relationship under HL, when the light intensity matched the growth environment (R^2^ = 0.38, p < 0.001; Fig. 4B). PRI showed the strongest overall association with maximum quantum yield (*F*v/*F*m) (R^2^ = 0.3, p <0.01; Fig. 4C), outperforming OVI which was the best predictor in the low GL experiment (Fig. 3B). PRI also displayed weak, positive associations with chlorophyll a, b and carotenoid contents under high GL (p < 0.05, Fig. 4A). Overall, these results indicate that PRI and NDVI capture variation in pigment composition and photosynthetic efficiency which are traits known to differ between light susceptible and light resilient duckweed varieties. Light stress susceptible *L. minuta* accessions could be separated from the better adapted *L. minor* and *L. japonica* accessions using PRI and NDVI for predicting *F*v/*F*m and *Φ*PSII (Fig. 4B-C). However, notably the highest R^2^ for any VI remained below 0.5 which is consistent with the low GL experiment, highlighting the limitations of VIs to be used as a solo proxy for vigour traits.

### 3.4. PLSR improves spectral prediction of pigments and photoprotection but not growth or biomass

To address the limitations of VIs for predicting duckweed traits, PLSR modelling was applied using spectral signatures ranging from 400-1100 nm, with the aim of improving prediction accuracy for novel crop traits. PLSR was used in the large duckweed cohort grown in low GL only where power was high enough to use training and validation. Whole-spectrum PLSR modelling substantially improved prediction of chlorophyll contents (Table 2, Fig. 5A) and significantly enhanced the prediction of baseline photoprotective traits compared with models using selected VIs (Fig. 5B). However, biomass and growth rate remained challenging to predict using whole spectral signatures.

**Fig. 5.**
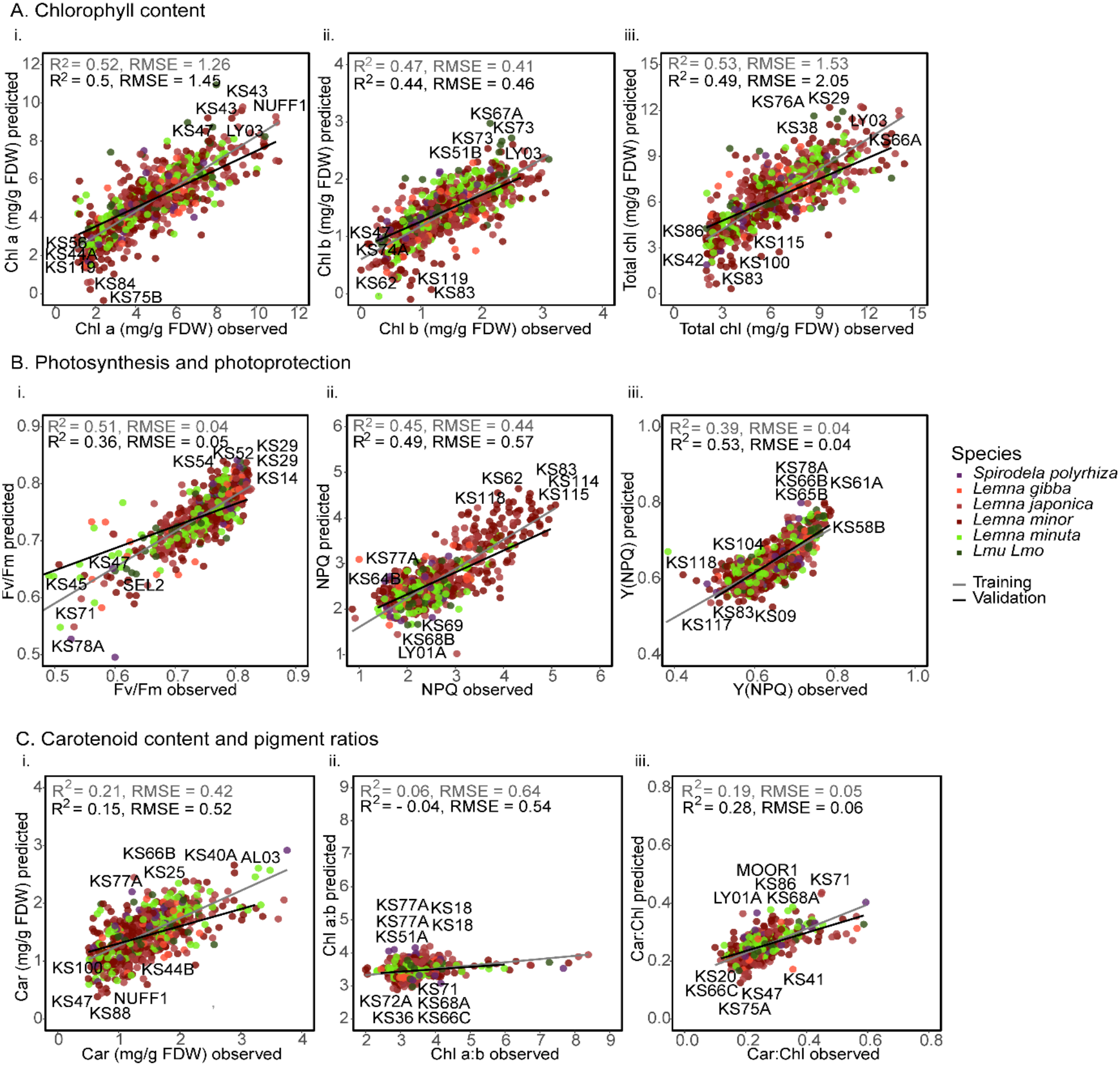
Physiological trait prediction using PLSR modelling for hyperspectral signatures of 133 duckweed accessions from low GL experiments. Observed measurements plotted against prediction derived from PLSR of hyperspectral data. **A.** Chlorophyll content. **B.** Photosynthetic and photoprotection traits. **C.** Carotenoid content and pigment ratios. Model training results are marked with grey sloped lines while validation results are indicated with black sloped lines. Top and bottom five performing accessions from the validation models are labelled in each panel.

**Table 2.** PLSR models of pigment, photosynthesis, photoprotection and growth traits using hyperspectral reflectance (400–1100 nm). Training (80%) and validation (20%) datasets were split by species. Model outputs include R^2^, RMSE = root mean square error, model bias error and *n* = sample sizes.

|  | **Training** |  |  |  | **Validation** |  |  |  |
|---|---|---|---|---|---|---|---|---|
| **Trait (units)** | **R<sup>2</sup>** | **RMSE** | **Model bias** | ***n*** | **R<sup>2</sup>** | **RMSE** | **Model bias** | ***n*** |
| Chl a (mg/g FDW) | 0.52 | 1.26 | -0.01 | 523 | 0.5 | 1.45 | 0.06 | 123 |
| Chl b (mg/g FDW) | 0.47 | 0.41 | -0.002 | 520 | 0.44 | 0.46 | 0.02 | 133 |
| Total chl (mg/g FDW) | 0.53 | 1.53 | -0.02 | 514 | 0.49 | 2.05 | -0.14 | 132 |
| *F*<sub>v</sub>/*F*<sub>m</sub> (-) | 0.51 | 0.04 | -0.0001 | 527 | 0.36 | 0.06 | 0.005 | 134 |
| NPQ VHL (-) | 0.45 | 0.44 | 0.001 | 527 | 0.49 | 0.57 | -0.04 | 134 |
| Y(NPQ) VHL (-) | 0.39 | 0.04 | 0.0003 | 527 | 0.53 | 0.04 | 0.0018 | 134 |
| Car (mg/g FDW) | 0.21 | 0.42 | -0.001 | 477 | 0.15 | 0.52 | -0.03 | 122 |
| Chl a:b (-) | 0.06 | 0.64 | 0.0002 | 509 | -0.04 | 0.54 | 0.03 | 131 |
| Car:Chl (-) | 0.19 | 0.05 | 0.0001 | 482 | 0.28 | 0.06 | 0.0006 | 123 |
| FW (g) | 0.31 | 337.9 | -2.11 | 171 | -0.09 | 457.9 | -26.04 | 45 |
| RGRlog area (day<sup>-1</sup>) | 0.06 | 0.62 | 0.01 | 167 | 0.15 | 0.63 | -0.13 | 44 |

PLSR modelling performed well for all chlorophyll content estimates: chlorophyll a, b and total chlorophyll, each achieving R^2^ ∼ 0.5, with a good alignment between the validation and training models (Fig. 5A). Modelling with PLSR improved chlorophyll prediction compared to using the GM index alone, and accessions with low and high chlorophyll contents showed good separation and proximity to validation fitting slopes (Fig. 5A). The spectral region with the highest predictive contribution was 550 nm as indicated by higher peaks in variable importance in projection (VIP) scores (Fig. 6A). A secondary peak for chlorophyll at the red edge between red and NIR around 700 nm was also consistently important (Fig. 6). In contrast, the early NIR region 750 – 900 nm which is used in key VI calculations such as NDVI and EVI, did not contribute to trait projections.

**Fig. 6.**
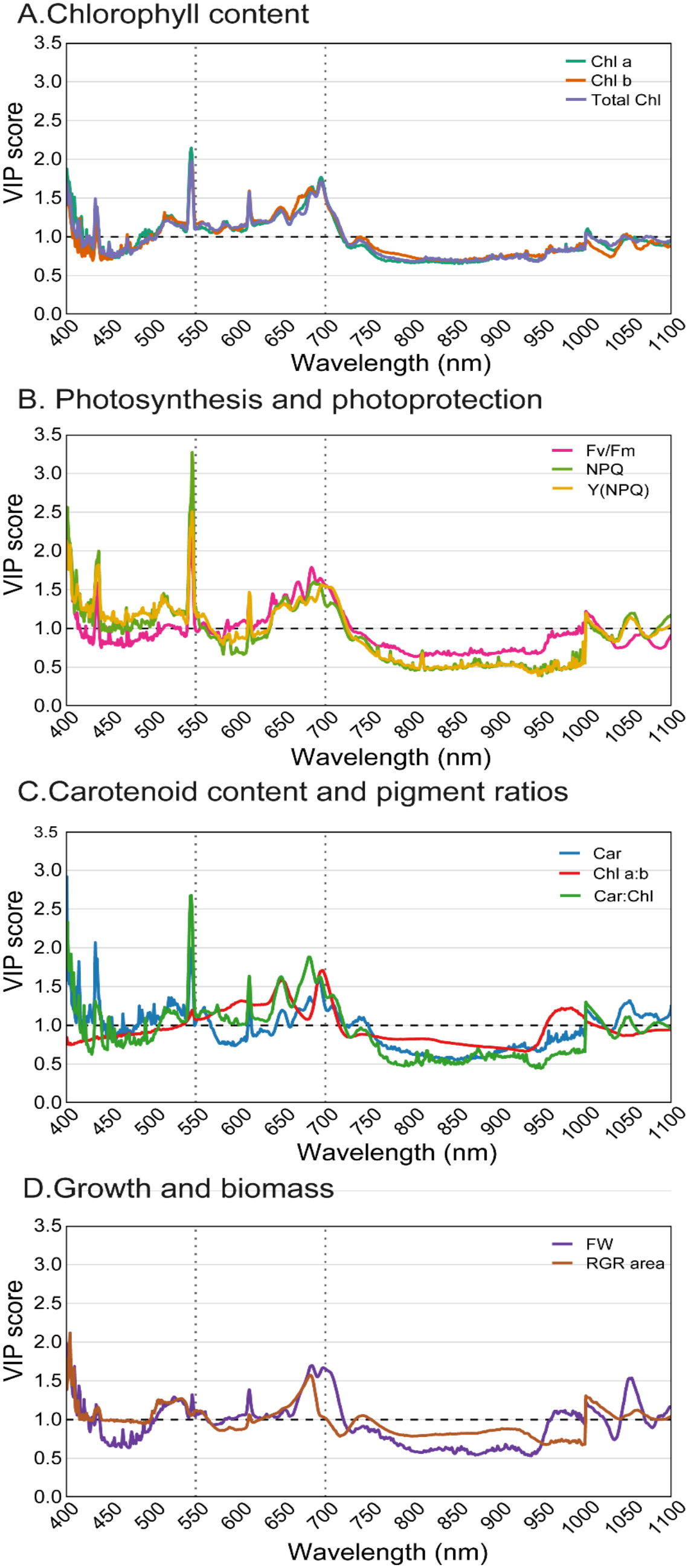
VIP scores from PLSR models reveal shared spectral regions contributing to the prediction of physiological traits in 133 duckweed accessions grown under low GL.. **A.** Chlorophyll content. **B.** Photosynthesis and photoprotection. **C.** Carotenoid content and pigment ratios. **D.** Growth and biomass. The dashed horizontal line indicates the VIP threshold for important predictors (VIP = 1). Dotted vertical lines highlight spectral regions near 550 and 700 nm, where multiple vigour-related traits showed strong predictive peaks.

For NPQ and regulated photoprotective dissipation (Y(NPQ)) at VHL, validation models outperformed the training models, yielding moderate R² values of 0.49 and 0.53, respectively (Table 2, Fig. 5B). The strongest VIP signal for photoprotection traits occurred at 550 nm, with Y(NPQ)photoprotection dissipation at VHL showing the highest VIP score of all modelled traits (Fig. 6). Notably, this region overlapped with chlorophyll and carotenoid contents. Although NPQ VHL did not correlate with any VI derived from NIR or visible spectra in either GL conditions, PLSR substantially improved prediction of photoprotective traits even in low GL conditions.. In contrast, validation performance for maximum quantum yield, *F*v/*F*m, was lower (R^2^ = 0.36) than the training results (R^2^ = 0.51, Table 2) indicating a poorer fit using PLSR. Divergence of slopes at the lower end of *F*v/*F*m values suggests that PLSR had limited predictive power for accessions with reduced photosynthetic efficiency (Fig. 5B).

PLSR modelling produced weak predictions for growth rate, biomass and pigment ratios (chlorophyll a:b and carotenoid:chlorophyll) (Table 2, Fig. 5). Notably, growth traits showed a high VIP score in the 650-700 nm region which overlapped with key wavelengths for predicting chlorophyll content and photoprotection traits NPQ and Y(NPQ) VHL. A distinctive peak between 1000-1050 nm was also identified for growth rate and biomass, corresponding to the NIR region characteristic of water absorption and commonly associated with vegetation moisture content, biomass and yield (Fig. 6). However, no trait (including growth rate and biomass) showed an association with water band indices such as NDWI (Fig. S1). For carotenoid content and pigment ratios, PLSR did not improve upon the low predictive power of VIs (Fig. 5C). Accessions were not well separated along model axes, especially for chlorophyll a:b where low and high performing accessions overlapped and predictions were especially poor at higher chl a:b ratios. Overall, these results suggest that hyperspectral signatures of duckweed biomass are effective for estimating total chlorophyll content but lack the sensitivity needed to reliably resolve variation in pigment ratios or to differentiate important growth and biomass trait capability of accessions.

## 4. Discussion

### 4.1. Limitations of hyperspectral reflectance information for biomass and growth prediction

Attainable biomass and growth rate are key traits to monitor for duckweed food production. Although accessions differed in their growth potential, hyperspectral VIs showed weak predictive power for biomass and growth rate. Red edge spectral regions around 680–710 nm associated with growth rates and biomass accumulation, yet PLSR models performed poorly. This contrasts with earlier duckweed studies, where VIs GM and OVI correlated with green area and fresh biomass during early growth in glasshouse conditions (Smith *et al*.; 2024c), and NDVI tracked *Lemna minor* biomass expansion in aquatic habitats (Gerardo & de Lima.; 2022). These studies measured biomass expansion coupled with water background reflectance compared to high density canopy endpoint measurements in this study. In fact, NDVI and EVI did not capture biomass variation sufficiently, and similarly low predictive power of NDVI was reported in other high density leafy systems such as mixed grasslands (R² = 0.4) (Ningthoujam *et al*.; 2025). In line with our results, red edge VIs including SR and EVI did not enhance biomass prediction accuracy for dense grassland (Fig. 3D, Fig. S1).

Under dense biomass conditions, leaf area and pigment content are maximized, maximizing high red absorbance for photosynthesis (low reflectance) and high NIR reflectance, resulting in saturation in calculated NDVI. Duckweed shows high NIR reflectance from dense mats (Ningthoujam *et al*.; 2025) particularly due to its floating adaptations including their waterproofing adaxial cuticle, intracellular air spaces for buoyancy and air-water interfaces (Cho *et al*.; 2008, Ziegler *et al*.; 2023). Reduced NIR reflectance occurs with lower vegetation coverage when water is visible between fronds in river habitats (Tian *et al*.; 2010). Therefore, in high density laboratory-standardised contexts, rather than NDVI capturing expanding leaf area and simultaneously reducing water reflectance as vegetation expands, signal sensitivity to growth dynamics is reduced and biomass prediction has an upper limit. NDVI readings above 0.6 are reported for healthy vegetation (Khdery *et al*.; 2026) but these values can be obtained even at low coverage (Murakami *et al*.; 2025), demonstrating that NDVI is sensitive to vegetation presence, but it does not increase linearly with biomass. Here, NDVI for accessions ranged between 0.3-0.75, showing variation in plant density and greenness under low GL growing conditions but a limit is reached to predict growth and biomass. Consequently, indices using R, red edge and NIR spectral regions such as NDVI, EVI, OVI and GM may be more informative to monitor greenness via pigments during active growth rather than under highly saturated conditions.

Treating duckweed like a standard arable crop also has limitations for yield prediction. Unlike conventional arable crops, duckweed remains vegetative and lacks the canopy transitions that can strengthen performance of VIs like NDVI and PRI for yield prediction. In crops, canopy organisation complexity and pigment composition have distinct vegetative and reproductive phases. For example, NDVI correlates moderately with chlorophyll content in the underutilised legume crop Bambara groundnut (*Vigna subterranea*) during vegetative growth (R² = 0.57) which improved during reproductive stages (R^2^ = 0.78) (Suhairi *et al*.; 2020), and VIs produced better yield predictions at pod filling (Jewan *et al*.; 2020). Similar patterns were found in cereals, where NDVI could strongly predict yield, biomass and light interception in vegetative stages and at peak canopy closure (R² = 0.75–0.87) (Lopresti *et al*.; 2015; Robles-Zazueta *et al*.; 2021; Kumar *et al*.; 2022; Pratiwi *et al*.; 2025). In contrast, duckweed forms a dense, flat and structurally uniform canopy with no definitive switch to reproductive and pod filling stages, meaning that hyperspectral VIs could not simply be applied to duckweed biomass in the same way as arable crops with growth stage shifts.

Taken together, this suggests that canopy saturation, aquatic floatation characteristics and limited phenological complexity could restrict the sensitivity of VIs to biomass detection or yield in duckweed. Despite these limitations, hyperspectral reflectance was suitable for monitoring or predicting pigment traits related to growth, such as chlorophyll content.

### 4.2. Spectral regions for prediction of pigment contents in duckweed

Chlorophyll content prediction was achievable using the GM index, with further improvement achieved through PLSR modelling. GM uses reflectance from green 550 nm and red edge 750 nm, whereas the PLSR model consistently highlighted 550 nm but the red edge region around 700 nm as key chlorophyll predictors, aligning with chlorophyll spectral regions in forest (Gitelson *et al*.; 2003) and wheat canopies (Wu *et al*.; 2008). As hypothesised, hyperspectral reflectance could identify duckweed varieties which maintained greener fronds in low input (light) environments and offers a robust non- destructive alternative to chlorophyll assays. To support the development of duckweed as a commercial crop, hyperspectral sensors could be integrated alongside existing RGB imaging systems in controlled- environment agriculture and vertical farming platforms, as has been demonstrated for salad crop production (Qin *et al*.; 2023). Spectral markers within the green and red edge regions could be used to monitor crop vigour, detect light stress and optimise nutrient management throughout production.

A remaining challenge is the separation of overlapping chlorophyll and carotenoid signals and to overcome limitations of VIs and PLSR for carotenoid prediction. Integration of hyperspectral reflectance and transmittance measurements, similar to approaches developed for broadleaf species like tobacco (Falcioni *et al.;* 2023b), may improve carotenoid quantification and isolate chlorophyll-specific spectral responses. Although implementing transmittance measurements is more challenging in small aquatic plants such as duckweed, combining reflectance, transmittance and chlorophyll fluorescence data could improve discrimination of pigment pools. Improved carotenoid detection would be particularly valuable for assessing both nutritional quality under low light cultivation and photoprotective responses under high irradiance. For example, high light finishing treatments have been used to enhance zeaxanthin accumulation in *Lemna* (López-Pozo *et al*.; 2024) and there remains opportunities for hyperspectral approaches to optimize pigments for nutritional applications.

### 4.3. Spectral regions for photosynthetic and photoprotection predictions in duckweed

Photoprotective traits such as NPQ and Y(NPQ) at VHL showed the strongest PLSR prediction overall. Although this modelling was performed for low GL duckweeds, there may be promise for PLSR to predict photoprotection capacity and efficiency in HL-grown duckweeds. In comparison, greenness VIs in both low and high GL lacked sensitivity to predict maximum NPQ VHL (Fig. S1–S2), indicating that PLSR models captured more relevant wavelengths associated to photoprotection which simple VIs missed. The spectral regions driving NPQ and Y(NPQ) VHL prediction, particularly the green peak around 550 nm and the red edge between 700–750 nm, were the same regions that drove chlorophyll and carotenoid models. This spectral overlap reflects the fundamental role of chlorophyll and carotenoid maintenance and xanthophyll cycle conversion underpinning dissipation as NPQ. The red edge spectral region also associated with growth and biomass prediction, support the linking of pigment composition and photoprotection to overall vigour in light environments. Consistent with this association, strong collinearity was observed among pigment contents, photosynthetic traits (ETR, *F*v/*F*m, ΦPSII) and greenness-based VIs (PRI, GM, GI, NDVI, OVI) (Fig. S1). Therefore, suggesting that reductions in pigment content and photosynthetic performance are accompanied by reduced vigour in low GL. Together, these findings highlight the potential of hyperspectral monitoring of pigments and photosynthetic traits to optimise duckweed production. By integrating hyperspectral cameras and tracking spectral markers of pigments and photoprotection in real time, light intensity and spectral recipes could be tailored to maximise biomass accumulation or enhance photoprotective pigments for food applications. This would mirror techniques used to enhance biomass of microgreens under high intensity blue light (Dmitriev *et al*;. 2026) or could be used to increase xanthophylls in *Lemna*.

Vegetation indices moderately associated with photosynthetic traits in high GL duckweed; NDVI showed a positive relationship with ΦPSII HL and VHL, while PRI was associated with both *F*v/*F*m and carotenoid:chlorophyll ratios. Carotenoid and specifically zeaxanthin levels are low in low GL but accumulation is strongly influenced by light intensity. Therefore, high GL promotes xanthophyll cycle de-epoxidation, increasing zeaxanthin production to dissipate excess absorbed energy through NPQ (Stewart *et al*.; 2021; Smith *et al*.; 2024a). As zeaxanthin absorbs light strongly at 531 nm, this contributes to the importance of the green peak and is used in calculating the PRI index in traditional crops. In duckweed, PRI was negative and narrow in range (0 to -0.03), which likely reflects variation in light stress, including both short-term xanthophyll cycle activity and longer-term changes in chlorophyll–carotenoid balance (Gamon *et al*.; 2015; Ogawa *et al*.; 2024).

Although NDVI and PRI captured some variation in photosynthesis and pigment dynamics in duckweed from high GL conditions, chlorophyll fluorescence remains essential for detecting immediate photosynthetic adjustment to light. For example, PRI responded slower to thermal stress than *F*v/*F*m in wheat (Cao *et al*.; 2019), but such a temporal delay is unknown in duckweed. Together with relatively low predictive performance of VIs for trait association in duckweed below R^2^ = 0.5, potential lack of sensitivity and possible temporal delay, this may limit independent use of hyperspectral reflectance to characterise duckweed productivity under fluctuating light environments. Therefore, hyperspectral reflectance cameras should be developed alongside fluorometers to compliment photosynthetic data and together with RGB cameras to track growth. This connected and automated approach could be used to maximize duckweed productivity in high irradiance or under niche light recipes (Wong *et al*.; 2020, Lanoue *et al*.; 2026). Furthermore, there remains opportunities for integrating hyperspectral information with photosynthetic efficiency through both light-dependent electron transport and CO₂ assimilation measurements to elucidate the mechanisms underpinning growth rate variation in duckweed (Rintoul *et al*.; 2026). To this end, more nuanced components of chlorophyll fluorescence kinetic parameters (e.g. Falcioni *et al.;* 2023a) could be described for duckweed to improve proxies relevant to food production under different light environments.

## 5. Conclusions

This study demonstrates that hyperspectral reflectance can capture meaningful physiological variation in duckweed, particularly for chlorophyll content and photoprotection traits, however, the predictive power for biomass and growth rate was limited. Furthermore, uncoupling specific chlorophyll and carotenoid content responses presented challenges using hyperspectral reflectance alone. The use of multivariate statistical approaches, such as PLSR, provides a strong framework for predicting chlorophyll and key photosynthetic traits, demonstrating the value of full spectrum modelling rather than simple VIs applied from other crops. Future work should compare additional machine learning approaches and integrate hyperspectral data with other measurements to elucidate physiological mechanisms driving duckweed growth variation.

Overall, this work establishes a framework for integrating hyperspectral information with vigour traits to accelerate the selection of high-performing duckweed varieties for sustainable food production, which could be expanded to include other duckweed species. Species-level differences identified here, most notably the consistently high biomass, chlorophyll and carotenoid content of *Lemna japonica*, highlight promising accessions for low-input food production systems (Smith *et al*.; 2024a). *Lemna japonica* is also supported for upscaling in high GL due to its enhanced carotenoid yields. In contrast, the putative hybrid *Lmu/Lmo* could not be distinguished from *L. minuta* by pigments, growth or photosynthetic traits, suggesting these accessions could be unified taxonomically as *Lemna minuta* (Smith *et al*.; 2025).

Finally, extending hyperspectral approaches to the remote prediction of nutrient composition represents an important next step for duckweed production systems. Chlorophyll content, for example, is closely tied not only to photosynthetic performance but also to nutritional components such as leaf N status (Yoder & Pettigrew-Crosby.; 1995). Hyperspectral monitoring has shown the potential to predict chlorophyll and nutrient status in other emerging crops (Wijewardane *et al*.; 2023), but this capability has yet to be established in duckweed despite substantial variation in nutrient composition in varieties grown under common conditions (Smith *et al*.; 2024b; 2025). Consequently, integrating hyperspectral monitoring of pigments, photoprotection and nutrient status could provide a foundation for precision duckweed cultivation, enabling biomass, nutritional quality and resource-use efficiency to be optimised simultaneously in future food production systems.

## 6. Supplementary data

The following supplementary data are available online.

**Table S1.** One hundred and thirty three accessions included in this study including species types, collection site coordinates, regions and native water light environment measurements from different seasons.

**Fig. S1.** Correlation matrix showing relationships between vegetation indices and pigment, photosynthetic and growth traits across 133 duckweed accessions grown under a common low light environment.

**Fig. S2.** Correlation matrix showing relationships between vegetation indices and pigment, photosynthetic, biomass and macronutrient traits across 24 duckweed accessions grown under a common high light environment.

## Acknowledgements

Thanks to Emma Jougla for assistance with chlorophyll extractions, Kade Marshall and Bethany Severn for assistance with fresh biomass. Thanks to Laura Briers for support with chlorophyll fluorescence and thanks to Hayley Newman for freeze-drying duckweed samples.

## Author contributions

KES: Conceptualization, Data curation, Formal analysis, Investigation, Methodology, Validation, Visualization, Writing – Original draft. CARZ: Conceptualization, Formal analysis, Investigation, Methodology, Software, Validation, Writing – Original draft. LC: Data curation, Methodology. MHY: Data curation, Methodology. LM: Formal analysis, Methodology, Software. EHM: Conceptualization, Funding acquisition, Project administration, Resources, Supervision. All authors contributed to writing – Review and editing the final draft.

## Data availability

The data from this study are available from the corresponding authors upon request.

