## Supplementary Figures 1 & 2 for "Illuminating duckweed: hyperspectral reflectance highlights a predictive ceiling for accurate determination of biomass"

**Supplementary Figure 1.** Correlation matrix showing relationships between vegetation indices and pigment, photosynthetic and growth traits across 133 duckweed accessions grown under a common low GL environment.

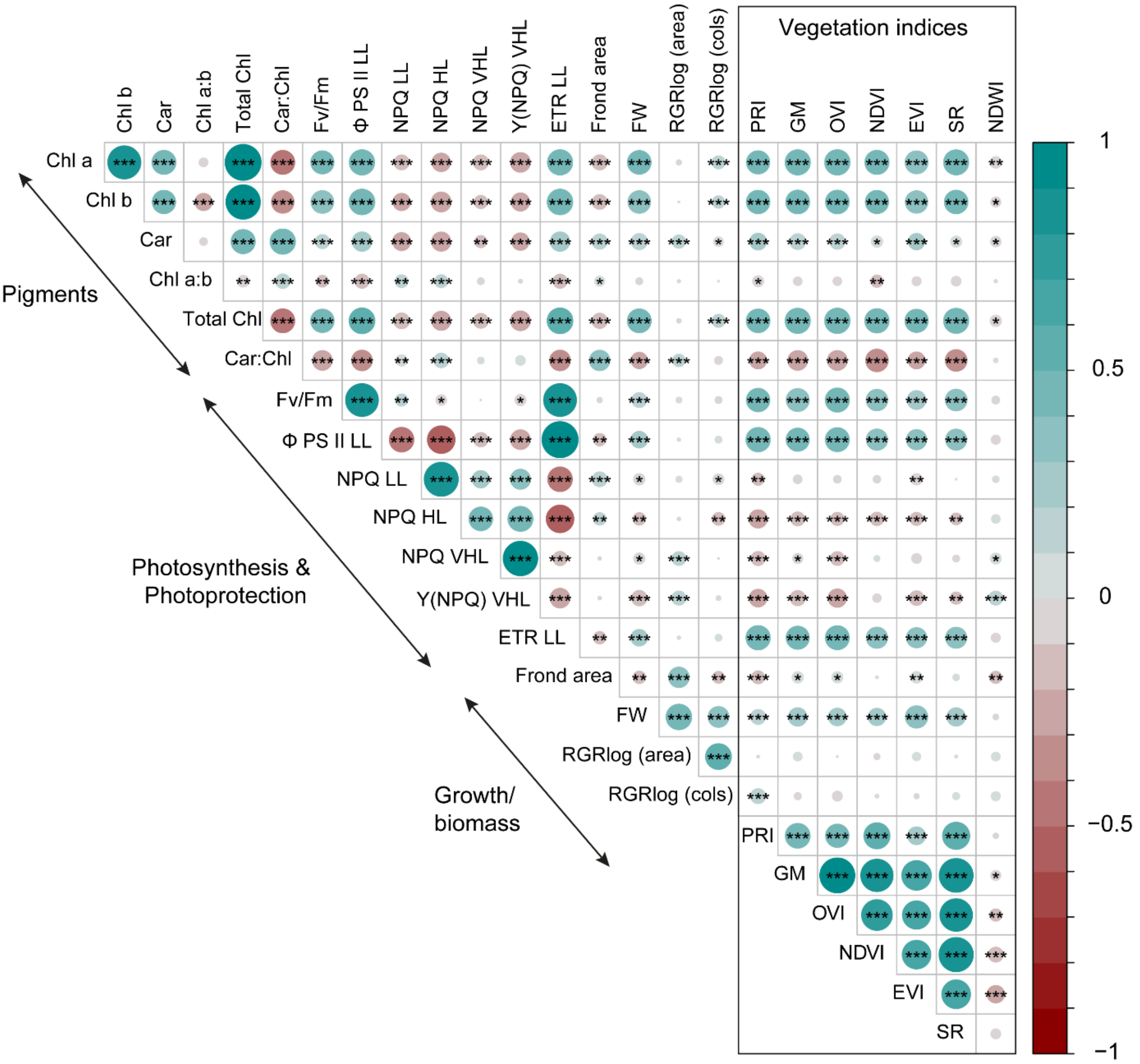


Circle size represents correlation coefficients, with blue indicating positive and red negative associations. Statistical significance is denoted with asterisks: P = < 0.05 (*), P = < 0.01 (**), and P = < 0.001 (***). Photosynthetic traits include maximum quantum efficiency of PSII ($F_v/F_m$,), operating efficiency of PSII at low light (ΦPSII LL), non-photochemical quenching under low, high and very high simulated light (NPQ LL, NPQ HL, NPQ VHL), regulated thermal dissipation at very high light Y(NPQ) VHL, electron transport rate under low simulated light (ETR LL) from chlorophyll fluorescence light response curves, frond area of initial colony at T0, fresh weight biomass (FW) and relative growth rates based on frond area and colony numbers from the log growth phase (RGRlog area, RGRlog col). Vegetation indices include photochemical reflectance index (PRI), green model (GM), optimal vegetation index (OVI), normalized difference vegetation index (NDVI), enhanced vegetation index (EVI), simple ratio (SR) and normalized difference water index (NDWI).

**Supplementary Figure 2.** Correlation matrix showing relationships between vegetation indices and pigment, photosynthetic traits, biomass and macronutrients across 24 duckweed accessions grown under a common high GL environment.

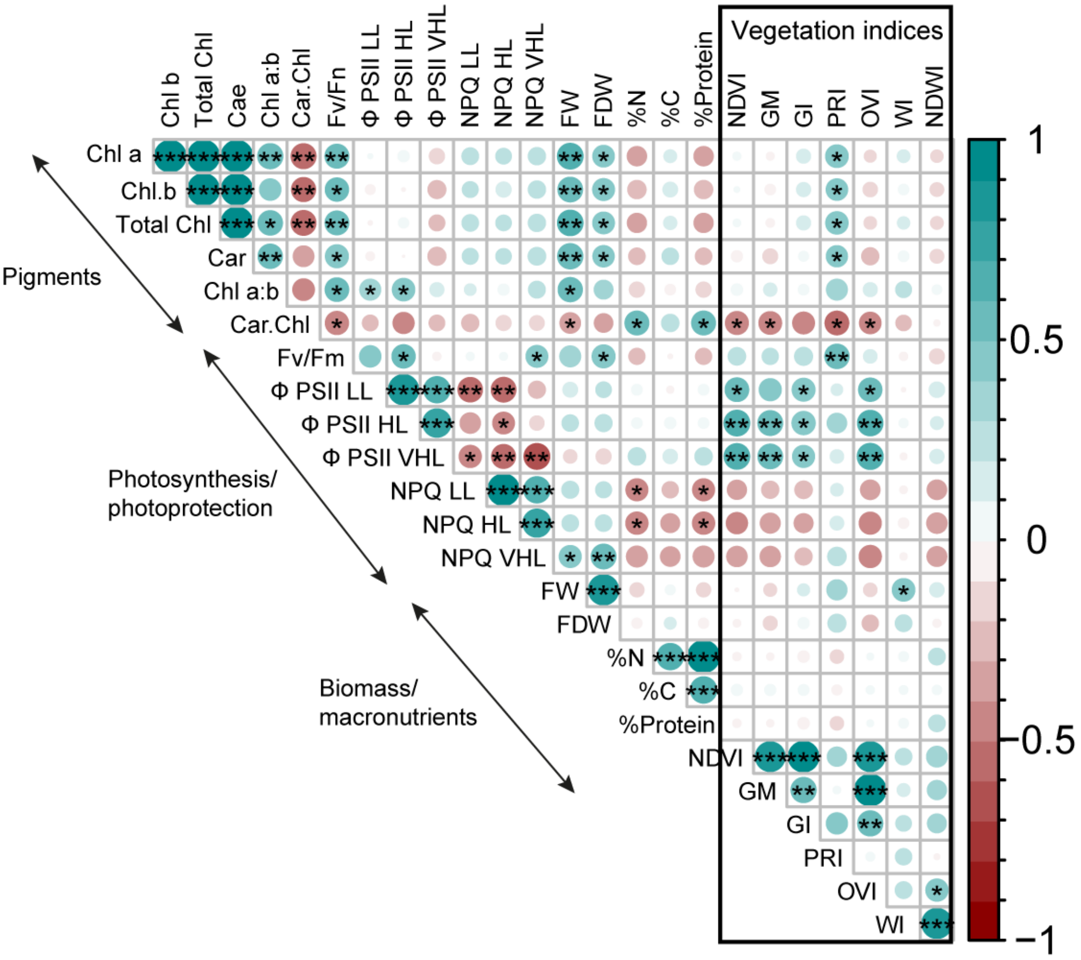


Circle size represents correlation coefficients, with blue indicating positive and red negative associations. Statistical significance is denoted with asterisks: P = < 0.05 (*), P = < 0.01 (**), and P = < 0.001 (***). Photosynthetic traits include: maximum quantum efficiency of PSII ($F_v/F_m$,), operating efficiency of PSII at low, high and very high light (ΦPSII LL, HL, VHL), non-photochemical quenching under simulated low, high and very high light (NPQ LL, NPQ HL, NPQ VHL) from light response curves. Fresh weight biomass (FW) and freeze dried weight (FDW), macronutrient contents as %C, %N and %Protein. Vegetation indices include normalized difference vegetation index (NDVI), green model (GM), green index (GI), photochemical reflectance index (PRI), optimal vegetation index (OVI), Water index (WI) and normalized difference water index (NDWI).
